# Heterozygous and generalist MxA super-restrictors overcome breadth-specificity tradeoffs in antiviral restriction

**DOI:** 10.1101/2024.10.10.617484

**Authors:** Rechel A. Geiger, Damini Khera, Jeannette L. Tenthorey, Georg Kochs, Laura Graf, Michael Emerman, Harmit S. Malik

**Author notes:** Correspondence should be addressed to: Harmit Singh Malik, 1100 Fairview Avenue N. A2-205, Seattle WA 98109.

## Abstract

Antiviral restriction factors such as MxA (myxovirus resistance protein A) inhibit a broad range of viruses. However, they face the challenge of maintaining this breadth as viruses evolve to escape their defense. Viral escape drives restriction factors to evolve rapidly, selecting for amino acid changes at their virus-binding interfaces to regain defense. How do restriction factors balance the breadth of antiviral functions against the need to evolve specificity against individual escaping viruses? We explored this question in human MxA, which uses its rapidly evolving loop L4 as the specificity determinant for *orthomyxoviruses* such as THOV and IAV. Previous combinatorial mutagenesis of rapidly evolving residues in human MxA loop L4 revealed variants with a ten-fold increase in potency against THOV. However, this strategy did not yield improved IAV restriction, suggesting a strong tradeoff between antiviral specificity and breadth. Here, using a modified combinatorial mutagenesis strategy, we find ‘super-restrictor’ MxA variants with over ten-fold enhanced restriction of the avian IAV strain H5N1 but reduced THOV restriction. Analysis of super-restrictor MxA variants reveals that the identity of residue 561 explains most of MxA’s breadth-specificity tradeoff in H5N1 versus THOV restriction. However, rare ‘generalist’ super-restrictors with enhanced restriction of both viruses allow MxA to overcome the breadth-specificity tradeoff. Finally, we show that a heterozygous combination of two ‘specialist’ super-restrictors, one against THOV and the other against IAV, enhances restriction against both viruses. Thus, two strategies enable restriction factors such as MxA to increase their restriction of diverse viruses to overcome breadth-specificity tradeoffs that may be pervasive in host-virus conflicts.

## Introduction

Pathogenic viruses pose persistent, ever-changing challenges to their hosts. To combat them, hosts encode various germline-encoded antiviral proteins, also called restriction factors, which form a critical component of innate immunity. Successful restriction by antiviral proteins often relies on their recognition and inhibition of viral targets in a cell-autonomous manner. However, these defenses must adapt to keep pace with viral target evolution and new viral challenges. Unlike adaptive immunity, which can adapt almost contemporaneously with viral evolution, the evolution of restriction factors is limited to rates of germline evolution^1^. Nonetheless, such evolutionary arms races drive rapid, recurrent amino acid changes (positive selection) at host-virus interaction interfaces, with hosts evolving to establish recognition of viral targets and viruses adapting to evade interaction with host restriction factors. Understanding the adaptive potential of rapid evolution in restriction factors is critical for understanding the biochemical and evolutionary constraints on viral restriction.

Because restriction factors are part of the genetically constrained innate immune system, they are often responsible for defending against a broad range of viruses with common molecular features^2^. MxA has one of the broadest antiviral ranges of any mammalian restriction factor; it can restrict multiple families of RNA and DNA viruses^2,3^. MxA is a dynamin-like, interferon-inducible GTPase whose antiviral activity, in most cases, depends on direct interactions with different viral targets^4–6^. MxA has been most extensively characterized for its restriction of *orthomyxoviruses* such as influenza A virus (IAV) and Thogotovirus (THOV)^7–12^. This antiviral activity requires GTP binding and hydrolysis by the globular G domain, oligomerization through sites in the stalk domain, and direct interactions with viral nucleoproteins (NP)^4,6,13–15^.

A growing body of evidence suggests that human MxA is a critical barrier to crossover events of animal-borne IAVs to humans^16–22^. Recently, it was shown that individuals infected with the avian IAV subtype H7N9 are more likely to have a dominant loss-of-function mutant MxA allele than healthy control groups^21^. Moreover, human-endemic IAV strains have acquired escape mutations in their NP proteins to resist human MxA restriction^16,23,24^, a prerequisite for continuous circulation in the human population^16,17,25^. For example, distinct mutations in the viral NP allowed the 1918 H1N1 pandemic IAV strain to evade human MxA and may have facilitated one of the most devastating IAV pandemics in humans^16^. In contrast, human MxA can restrict avian-derived influenza viruses such as the waterfowl-endemic H5N1 IAV, which is pathogenic in human individuals but has not acquired the capability for human-to-human transmission^23^. However, the recent rampant spread of an avian H5N1 IAV strain in domestic cattle populations in the US has raised renewed concerns about the possibility of zoonotic spillover events^26–28^. To better understand the ongoing arms race between MxA and human pathogenic *orthomyxoviruses*, we analyzed the evolutionary constraints that shape MxA restriction of *orthomyxoviruses*.

We previously identified the unstructured loop L4 of MxA, especially an aromatic residue (phenylalanine, tyrosine, or tryptophan) at amino acid position 561, as a critical determinant of THOV and IAV NP binding and restriction^29^. Furthermore, using combinatorial mutagenesis of five rapidly evolving residues in loop L4, we identified super-restrictor human MxA variants that could augment wildtype human MxA’s (hereafter referred to as wtMxA) already potent restriction of THOV^30^. In some cases, altering only a few positively selected residues in MxA L4 led to a ten-fold increase in potency. However, increased THOV restriction correlated with loss of restriction of the H5N1 strain of IAV (hereafter referred to as H5N1). These findings suggested a ‘breadth-specificity’ tradeoff in MxA restriction of *orthomyxoviruses*, wherein variants with increased potency against one virus lost specificity against another^30^.

Understanding the basis of such breadth-specificity tradeoffs and identifying strategies by which restriction factors like MxA can overcome them is critical to understanding their role in the human species’ barrier to potentially zoonotic viruses. In the present study, we modified our combinatorial mutagenesis strategy to identify human MxA super-restrictor variants with over 10-fold higher restriction of H5N1 IAV than wtMxA. By comparing H5N1-specific versus THOV-specific super-restrictors, we showed that the identity of the aromatic residue at residue 561 explained most of the breadth-specificity tradeoff; phenylalanine or tyrosine favored THOV restriction, whereas tryptophan favored H5N1 restriction. Despite this strong bias, we were nevertheless able to identify rare ‘generalist’ super-restrictor MxA variants that can simultaneously restrict both THOV and H5N1 more potently than wtMxA, thereby providing an intrinsic means to overcome the breadth-specificity tradeoff. Moreover, combining two ‘specialist’ restrictors of THOV and H5N1 in heterozygous combinations provided an extrinsic means to improve potency against both viruses, allowing the host to benefit from each allele without suffering from dominant-negative interference. Our study reveals the basis of breadth-specificity tradeoffs that constrain the evolution of host restriction factors like MxA against multiple pathogenic viruses and two strategies that may help to overcome them.

## Results

### Combinatorial mutagenesis of loop L4 reveals MxA variants with enhanced H5N1 restriction

To understand how an antiviral protein can improve defense against a specific viral target through changes to rapidly evolving residues, we previously carried out two combinatorial mutagenesis screens to identify potential MxA super-restrictors with greater than ten-fold improved restriction of THOV^30^. However, the MxA variants with the highest levels of THOV restriction were impaired in their restriction activity against H5N1 IAV, revealing a breadth-specificity tradeoff. Here, we sought to identify H5N1 super-restrictor variants of MxA and to identify pathways through which the host might evolve around this breadth-specificity tradeoff.

Like with THOV restriction^30^, we first confirmed that non-aromatic residues at residue 561 were incompatible with H5N1 restriction (Fig. S1, Supplementary Table S1). Our previous study revealed that W561 variants maintained potency against H5N1 despite losing THOV restriction^30^. Therefore, we hypothesized that we could select H5N1 super-restrictors more successfully if we allowed residue 561 to be either F, Y, or W instead of restricting it to F alone^30^. We generated a new combinatorial mutagenesis library in which residue 561 sampled only the aromatic amino acids while allowing the other four rapidly evolving residues (540, 564, 566, and 567) to encode any amino acid (see Methods) (Fig. 1A). We randomly selected nearly 200 unique MxA variants from this combinatorial mutagenesis library, only discarding variants with stop codons introduced during NNS mutagenesis or variants with missense mutations outside loop L4. We then tested these MxA variants for their ability to restrict H5N1 (A/Vietnam/1203/04) using a previously described minireplicon assay^24^ (see Methods), which assesses the effects of MxA restriction on viral transcription and genome replication^24,29–31^. Briefly, we co-expressed all components of the viral ribonucleoprotein (vRNP), including the polymerase complex, the nucleoprotein (NP), an artificial, viral RNA genome segment encoding a reporter firefly luciferase gene that is transcribed and replicated by the viral polymerase in an NP-dependent manner, and a transfection control *Renilla* luciferase reporter. NP-dependent polymerase activity is measured as a ratio of firefly-to-*Renilla* luciferase. We calculated the fold restriction as the decrease of luciferase activity of the H5N1 minireplicon in the presence of co-transfected MxA. We used catalytically dead human MxA (T103A) as a negative control for restriction and multiple replicates of wtMxA to define the wildtype range of H5N1 restriction.

**Figure 1:**
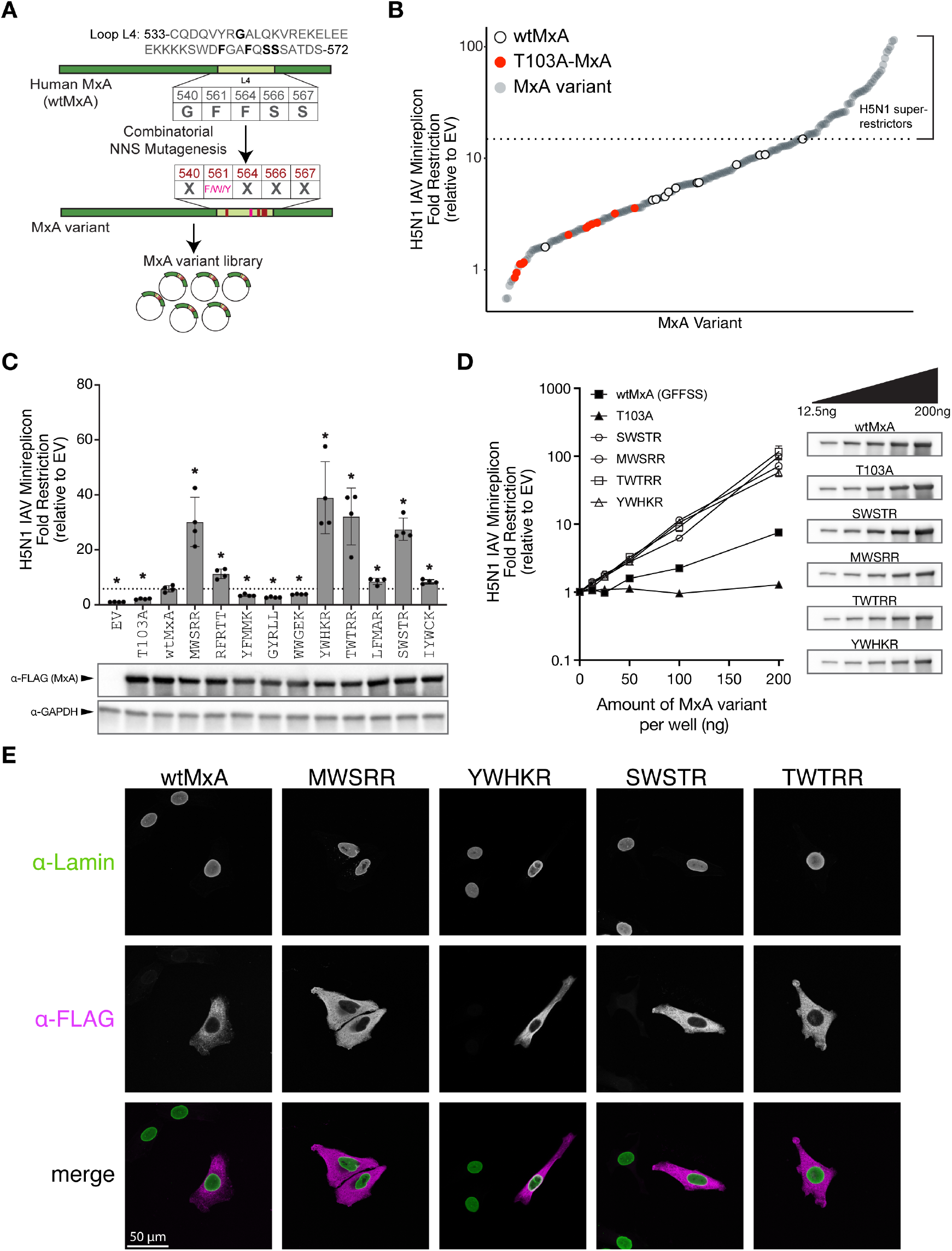
Combinatorial mutagenesis of human MxA identifies H5N1 super-restrictors. **(A)** An Mx variant library was constructed using combinatorial mutagenesis. Amino acid residues 540, 564, 566, and 567 (top, bolded) from loop L4 of wildtype human MxA were allowed to mutate to any amino acid residue using NNS mutagenesis, whereas residue 561 was varied to encode either F, W, or Y. **(B)** Restriction profile of 194 unique MxA combinatorial variants in L4 (gray circles) against the H5N1 minireplicon activity relative to empty vector control (on a log_10_ scale). For comparison, we also include multiple replicates of a catalytically inactive GTPase MxA variant, T103A (red circles), and several independent replicates of wtMxA (open circles) to define the wildtype range of H5N1 restriction. All analyses were performed in biological triplicates, and average restriction is represented. We used the highest level of wtMxA restriction to define the threshold level of H5N1 restriction (dotted line), above which the variants were tentatively classified as ‘super-restrictors.’ **(C)** Ten of the top super-restrictors were re-evaluated for H5N1 restriction as a fold difference to the empty vector control using an independently performed minireplicon assay restriction in biological quadruplicates (on a linear scale). Variants are identified based on their amino acid identities at the five variable sites in MxA L4, *i*.*e*., 540, 561, 564, 566, and 567. We used unpaired Welch’s t-tests between each variant and wtMxA to evaluate statistical significance (*p-value < 0.05). Protein expression levels of each variant were monitored by Western blotting. **(D)** Four validated super-restrictors were tested for their H5N1 restriction relative to empty vector (on a log_10_ scale) at increasing doses of MxA expression. The total amount of transfected DNA was equalized in all experimental conditions by supplementing empty vector plasmid DNA. Protein expression levels of each variant at each dosage were monitored by Western blotting. **(E)** Immunofluorescence analysis of the sub-cellular localization of wtMxA and four validated MxA super-restrictor variants using FLAG epitope-tagging in transiently transfected HeLa cells, which were also stained for lamin (which localizes to the nuclear membrane).

We found that wtMxA has a relatively modest restriction of H5N1 (average restriction of 6.85-fold relative to empty vector control, Fig. 1B), consistent with previous studies^16,30^. However, 51 out of 194 MxA combinatorial variants showed higher H5N1 restriction than the highest level of restriction observed among multiple wtMxA replicates – in some cases up to 15-fold (we refer to variants with this improved restriction as ‘super-restrictors’) (Fig. 1B, Supplementary Table S2). To validate our hits, we retested the H5N1 restriction activity of ten of the top super-restrictor variants obtained in our initial screen using an independently performed minireplicon assay (Fig. 1C, Supplementary Table S3). This reanalysis reconfirmed the improved H5N1 restriction activity of 7 out of 10 MxA variants. All variants tested express MxA protein at similar levels (Fig. 1C). Therefore, increased restriction of the top super-restrictors does not result from higher expression or stability. We retested the dose-dependent restriction for four validated super-restrictors – SWSTR, MWSRR, TWTRR, and YWHKR, where each letter refers to the positions of the positively selected residues in L4 of human MxA – 540, 561, 564, 566, and 567, respectively. The titration confirmed the enhanced H5N1 restriction activity of the super-restrictor variants compared to wtMxA at multiple levels of plasmid input (Fig. 1D, Supplementary Table S4).

Since nuclear-localized MxA has enhanced IAV restriction^32–34^, we tested whether altered subcellular localization could underlie the enhanced activity of these top four variants. We visualized FLAG-tagged versions of these MxA variants in HeLa cells (see Methods). We found that the super-restrictors localized diffusely to the cytoplasm, like wtMxA (Fig. 1E). Moreover, in all cases, appending an SV40 NLS to their N-termini (Methods) could drive their nuclear localization (Fig. S2A) and enhance H5N1 restriction (Fig. S2B, Supplementary Table S5) without altering their expression levels. In contrast, the nuclear re-localization of a non-super-restrictor (WWGEK) was insufficient to enhance its potency over NLS-wtMxA. Therefore, we conclude that the H5N1 MxA super-restrictors act in their native cytoplasmic location without altered subcellular localization. These findings confirm that the sequence space of the rapidly evolving residues in loop L4 includes MxA variants with super-restriction of diverse *orthomyxoviruses*.

### Positive epistasis between L4 residues underlies H5N1 super-restriction in MxA variants

Next, we aimed to understand the amino acid patterns underlying H5N1 super-restriction. Using a DiffLogo plot, we compared the frequencies of MxA L4 residues found in H5N1 super-restrictors from Figure 1B versus all other tested variants (Fig. 2A). This comparison revealed a clear preference for tryptophan at position 561 (W561) among H5N1 super-restrictors, whereas Y561 was strongly disfavored. F561 (the residue in wtMxA) showed no strong preference, which is why it is barely visible in the DiffLogo plot. This pattern is even more apparent when the H5N1 restriction activities of all tested MxA variants are grouped based on residue 561 (Fig. 2B, Supplementary Table S1). These findings sharply contrast with our previous analysis of THOV super-restrictors, which favored either Y561 or F561 and disfavored W561^30^. The DiffLogo plot also highlighted a striking preference for positively charged residues (arginine [R] or lysine [K]) at positions 564, 566, and 567 (Fig. 2A).

**Figure 2:**
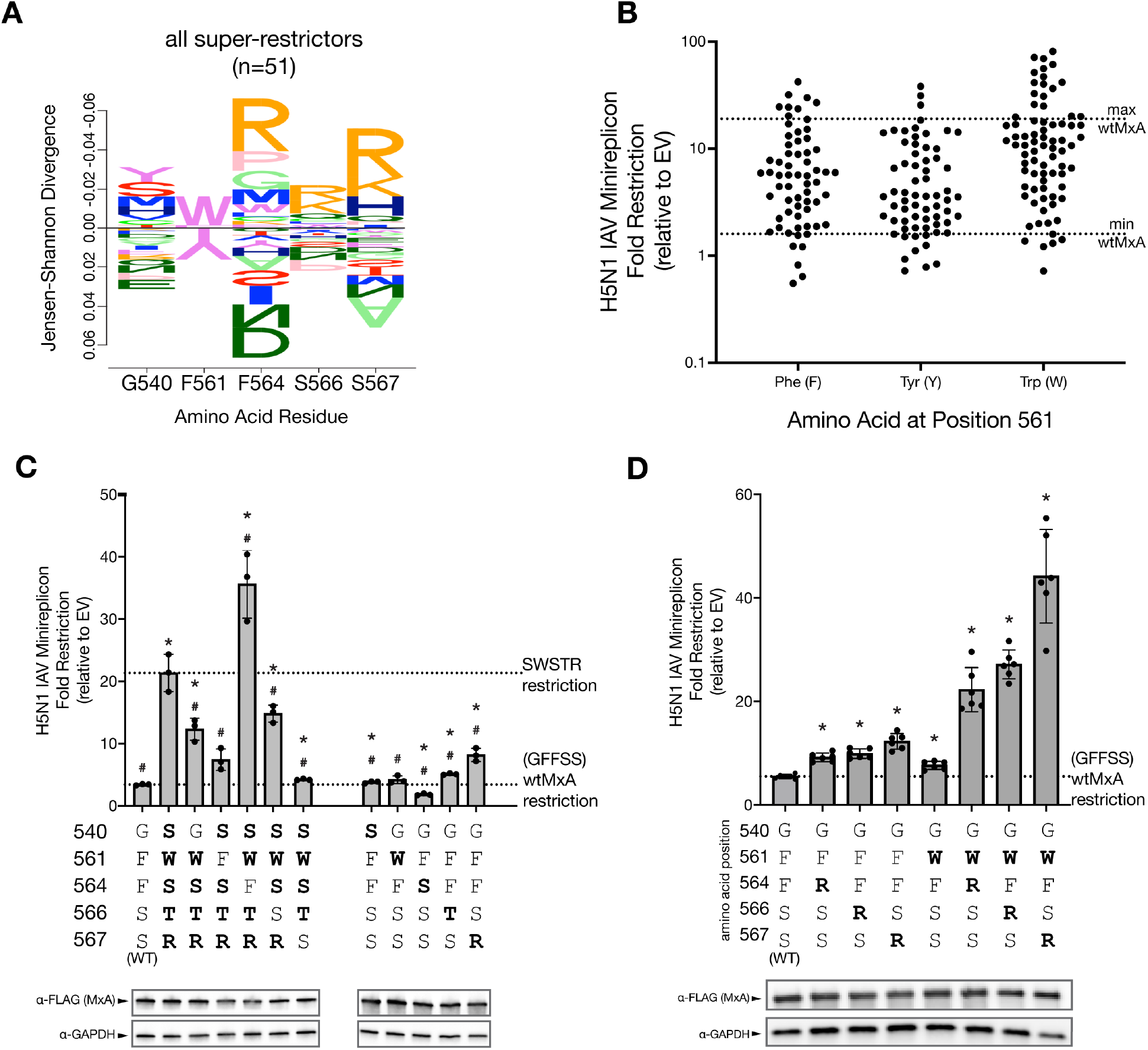
Necessity and sufficiency of L4 residues reveal positive epistasis underlies MxA super-restriction of H5N1. **(A)** DiffLogo plot compares residue frequencies among super-restrictors (variants with restriction greater than maximal wtMxA restriction) relative to all tested variants, indicating the amino acid preferences at each position for H5N1 super-restriction. Bar height is proportional to Jensen-Shannon divergence per site, and letter height is proportional to frequency. The wtMxA residues (GFFSS) are indicated at the bottom. **(B)** Dot plot showing the restriction profiles (as fold restriction relative to empty vector, log_10_ scale) of all 194 MxA variants, sorted according to the amino acid identity at residue 561. The dotted lines indicate the highest and lowest restriction levels of wtMxA (taken from Figure 1B). **(C)** We analyzed the contribution of each residue of super-restrictor variant SWSTR in conferring increased anti-H5N1 activity by reverting each position to the wtMxA residue, GFFSS (left). We also tested the sufficiency of individual changes in L4 residues from the SWSTR variant to confer increased H5N1 restriction to wtMxA (right). Protein expression levels of each variant were monitored by Western blotting. **(D)** We analyzed the ability of dual changes in L4 residues – F561W combined with either 564R, 566R, or 567R – to confer increased H5N1 super-restriction to wtMxA. Protein expression levels of each variant were monitored by Western blotting. The restriction is reported as a fold-change relative to an empty vector on a linear scale. For **(C)**, we performed unpaired Welch’s t-tests between each variant and wtMxA (asterisks, p-value < 0.05*) or SWSTR (hashes, p-value < 0.05 #). For **(D)**, we performed unpaired Welch’s t-tests between each mean and the wtMxA mean. Significant differences (p-value < 0.05) are noted as *.

Next, we investigated the contribution of the single residues at the five variable positions of a selected super-restrictor for enhanced H5N1 restriction. The SWSTR variant was one of the strongest H5N1 super-restrictors in our initial and revalidated screen (Fig. 1), with ∼5-fold higher H5N1 restriction than wtMxA (GFFSS) (Fig. 2C, far left). We found that individually reverting residues 540, 561, 566, and 567 in SWSTR to wtMxA (*i*.*e*., S540G, W561F, T566S, or R567S) led to a significant reduction in H5N1 restriction (Fig. 2C, Supplementary Table S6). W561F and R567S reversions showed the most dramatic decrease in restriction, consistent with the preference for both 561W and 567R among super-restrictor variants (Fig. 2A). In contrast, we found that reversion of residue 564 (*i*.*e*., S564F) increased H5N1 restriction, consistent with 564S being disfavored among super-restrictors (Fig. 2A). The resulting SWFTR variant is 10-fold better than wtMxA at restricting H5N1 (Fig. 2C, Supplementary Table S6). Thus, multiple residues in the SWSTR variant, most notably W561 and R567, are necessary for the increased H5N1 activity of super-restrictor variants.

We also tested which residues of SWSTR were sufficient to confer H5N1 super-restriction to wtMxA (Fig. 2C). Several single residue changes from SWSTR into the wtMxA backbone led to statistically significant increases in restriction. However, these increases were modest, with S567R providing the largest 2.4-fold increase of H5N1 restriction over wtMxA (Fig. 2C, right). Thus, we conclude that robust H5N1 super-restriction by SWSTR requires multiple changes from wtMxA. Therefore, we tested whether two changes might be sufficient to confer SWSTR-like levels of super-restriction onto wtMxA (Fig. 2D). Since the DiffLogo plot indicated a preference for W561 and basic residues at positions 564, 566, and 567 among H5N1 super-restrictors (Fig. 2A), we introduced W561 in conjunction with either 564R, 566R, or 567R in wtMxA. We found that each of these combinations conferred H5N1 super-restriction levels that were higher than the sum of individual mutations (Fig. 2D), Supplementary Table S7), reiterating the importance of the W561 residue and the interchangeability of the arginine residue at either residue 564, 566, or 567. These findings imply that positive epistatic interactions among at least two residues of loop L4 are required to achieve H5N1 super-restriction, as with THOV super-restriction^30^.

### ‘Generalist’ MxA variants overcome breadth-specificity tradeoffs in viral restriction

Our previous study revealed that increased THOV restriction often weakened H5N1 restriction^30^. We investigated this breadth-specificity tradeoff in more detail, aided by our identification of novel H5N1 super-restrictors. We selected 52 MxA combinatorial variants from our screen based on their H5N1 restriction (Fig. 1B): 42 super-restrictor variants with better than wtMxA restriction, five variants with equivalent restriction as wtMxA, and five non-restrictors with lower than wtMxA activity. We assayed all 52 MxA variants for their ability to restrict THOV (x-axis) and re-assayed them against H5N1 (y-axis) using minireplicon assays (Fig. 3A, Supplementary Table S8). By

**Figure 3:**
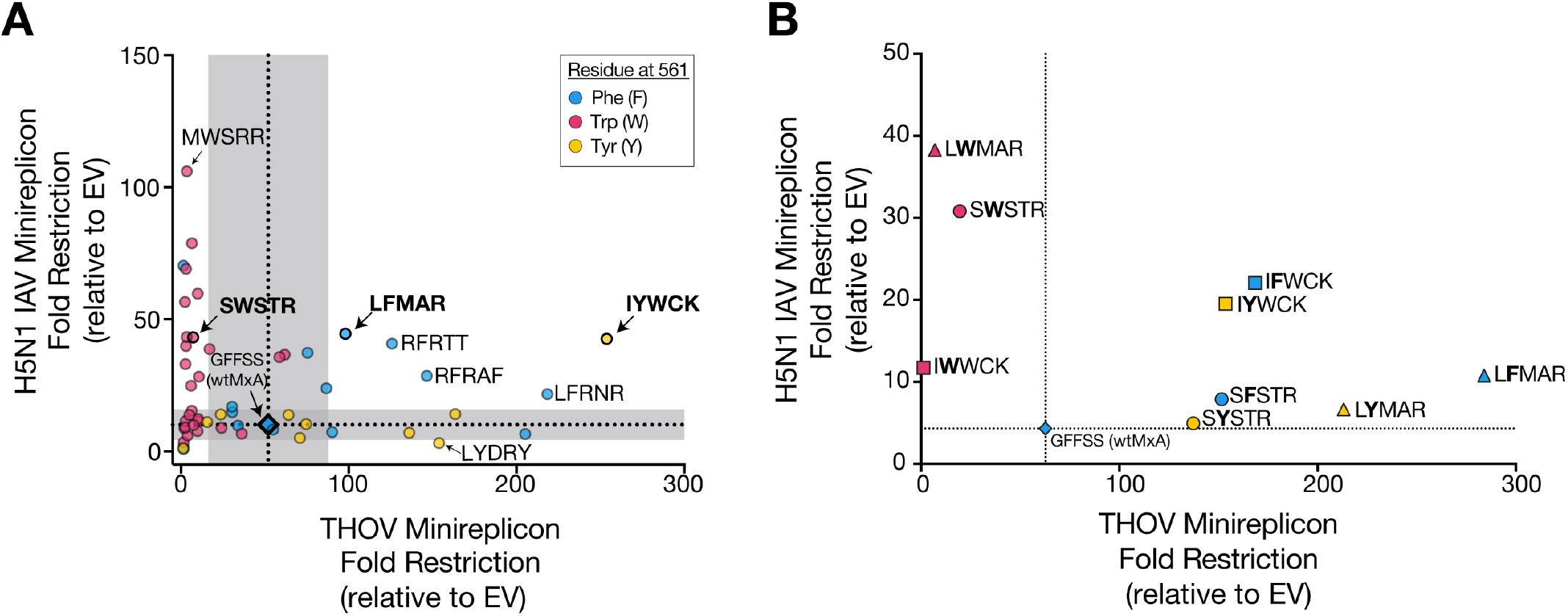
‘Generalist’ MxA variants can restrict both H5N1 and THOV. **(A)** We screened 52 MxA variants for antiviral activity in H5N1 and THOV minireplicon assays. Each point represents an average of three independent replicates. The mean wild-type restriction of H5N1 and THOV are represented by horizontal and vertical dotted lines, respectively, with gray shaded areas representing two standard deviations among three replicate wtMxA measurements. We classified MxA variants based on residue 561 – either phenylalanine (F, cyan dot), tyrosine (Y, yellow dot), or tryptophan (W, magenta dot). **(B)** We replaced the amino acid at residue 561 in selected super-restrictors with other aromatic residues. All points represent the average of three replicate experiments. X- and y-axes in linear scale.

comparing their activity to the range of wtMxA values (using two standard deviations above mean wtMxA activity as a threshold), we classified variants as H5N1 specialist super-restrictors (above the horizontal shaded region, Fig. 3A) or THOV super-restrictors (right of the vertical shaded region, Fig. 3A).

Consistent with a breadth-versus-specificity trade-off, most MxA super-restrictors are ‘specialists,’ *i*.*e*., they have increased H5N1 but not THOV restriction or increased THOV but not H5N1 restriction. Seventeen of 22 H5N1 specialist super-restrictors encode W561 (Fig. 3A, magenta circles). In contrast, none of the THOV specialist super-restrictors encoded W561; instead, they encoded F561 or Y561 (Fig 3A, blue and yellow circles). To further test the role of residue 561 on H5N1 versus THOV restriction, we tested the effects of swapping this residue with the other aromatic amino acids for three MxA super-restrictors containing either W561 (SWSTR), F561 (LFMAR), or Y561 (IYWCK) (bold, Fig. 3A, Supplementary Table S8) on their THOV or H5N1 restriction activity. For all three variants, we found that F561 and Y561 in all three variants produced robust THOV restriction, whereas W561 resulted in a dramatic loss of THOV restriction (Fig. 3B), confirming earlier findings that W561 may be incompatible with THOV restriction^30^ (Fig. 3A). Furthermore, we found a clear hierarchy (W561 > F561 >Y561) in H5N1 restriction for the SWSTR and LFMAR backgrounds (Fig. 3B), consistent with our DiffLogo analyses (Fig. 2A). Together, these results bolster our previous findings. Despite a few exceptions, we conclude that most of the breadth-specificity trade-off in THOV versus H5N1 super-restriction by MxA may stem from the identity of the aromatic amino acid found at MxA’s critical residue 561.

Unexpectedly, our analyses also revealed five ‘generalist’ super-restrictor MxA variants, which had much improved (greater than two standard deviations above mean wtMxA) restriction against both THOV and H5N1 (circles above and to the right of gray shaded areas, Fig. 3A). Intriguingly, four of these MxA variants encoded F561, while one encodes Y561. The IYWCK generalist super-restrictors also had an atypical sequence preference for H5N1 restriction, with IYWCK and IFWCK outperforming IWWCK (Fig. 3B, Supplementary Table S9). Although generalist super-restrictors constitute ∼10% of the 52 MxA variants tested for both viruses, this is likely an overestimate of the frequency of generalist super-restrictors among MxA combinatorial variants since we biased our selection towards H5N1 super-restrictors. Indeed, our previous analysis of 24 THOV super-restrictors found that most of them lost H5N1 restriction^30^. Our findings suggest that MxA’s breadth-specificity trade-off in H5N1 and THOV restriction is not insurmountable; generalist super-restrictor variants can simultaneously improve their intrinsic restriction of multiple divergent *orthomyxoviruses* such as THOV and H5N1.

### Heterozygous MxA ‘specialist’ variants combine to yield ‘generalist’ super-restriction

Generalist super-restrictor variants are rare in the evolutionary landscape compared to specialist super-restrictors (Fig. 3), whose enhanced antiviral activity can be achieved by just two amino acid changes (Fig. 2). We hypothesized that generalist super-restriction might still be achieved by combining two distinct specialist super-restrictors as heterozygous alleles. Two observations inspired this hypothesis. First, previous studies have shown that loss-of-function variants found in the human population can have a dominant-negative effect on wtMxA restriction of IAV, by ‘poisoning’ MxA oligomers required for antiviral activity^21,35^. Second, recurrent signatures of positive selection in MxA^29^ suggest that any *de novo* gain-of-function MxA variant must have been able to provide increased function even as a heterozygous allele with wtMxA.

To test this hypothesis, we first investigated whether a super-restrictor MxA variant could provide enhanced restriction in the presence of wtMxA, mimicking its origin as a heterozygous allele. We selected two H5N1 specialists (MWSRR, SWSTR), two THOV specialists (QFAYS, LYDRY), and one generalist (IYWCK). We re-tested them against H5N1 and THOV to confirm their specialist and generalist restriction activities (Figure S3, Supplementary Tables S10, S11). We then tested the H5N1 and THOV restriction activity of each MxA super-restrictor variant in a 1:1 ratio with wtMxA, mimicking equal amounts of heterozygous alleles (Fig. 4A-B, Supplementary Tables S12, S13). As dosage controls, we tested our wtMxA in a 1:1 ratio with either an empty vector (1X wtMxA) or another wtMxA three other variants in a 1:1 ratio with wtMxA for restriction: catalytically inactive MxA (T103A) previously reported to be dominant-negative to wtMxA^36^, an oligomerization-defective MxA variant (M527D)^5,15^, and a dominant-negative MxA variant identified in a human patient (L542S)^21^.

**Figure 4:**
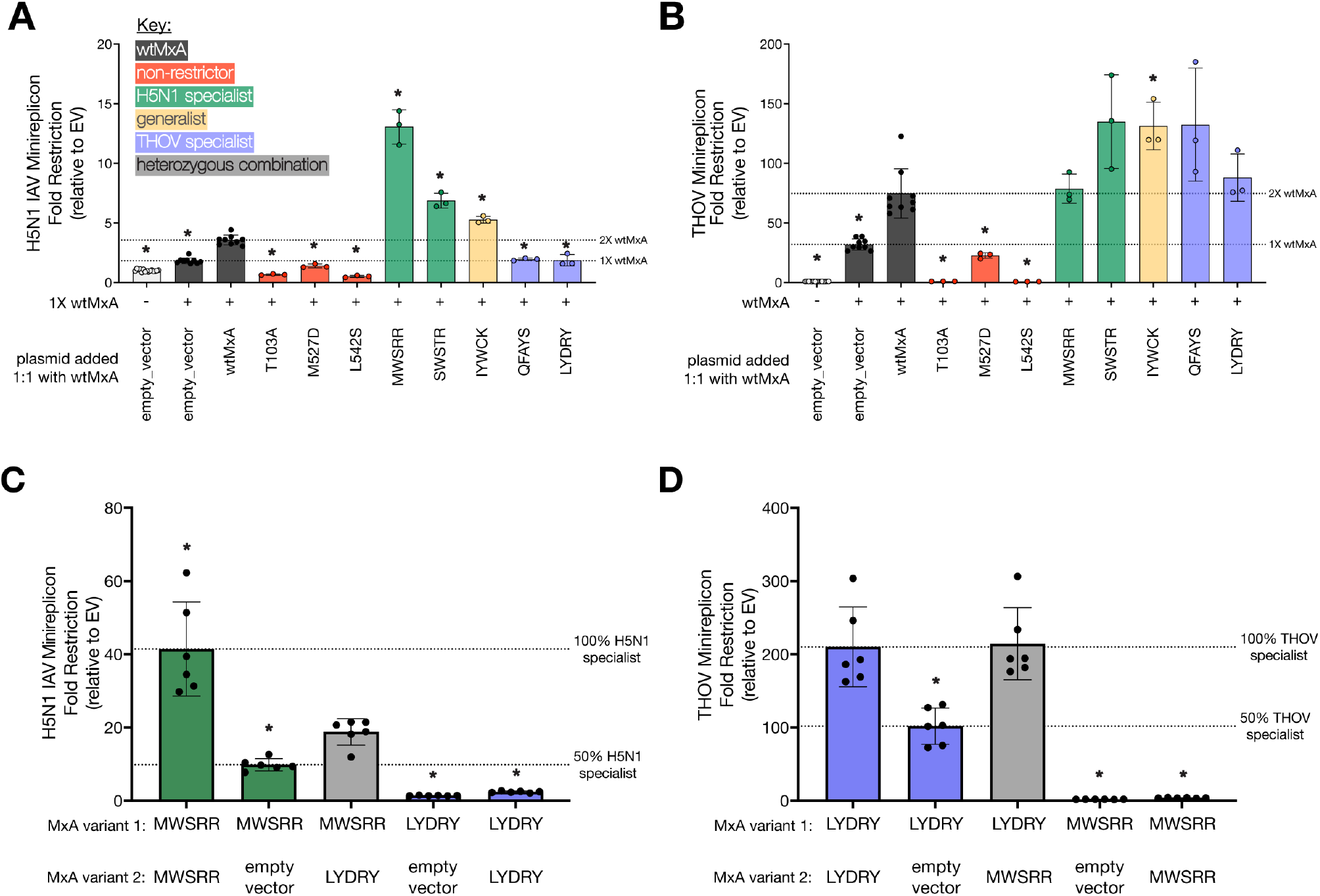
Heterozygous MxA ‘specialist’ variants combine to yield ‘generalist’ super-restriction of H5N1 and THOV. **(A)** Using minireplicon assays, we tested 1X wtMxA (100ng per well for H5N1 assay and 50ng per well for THOV assay) and equimolar ratios of different MxA variants, including wtMxA, for **(A)** H5N1 restriction and **(B)** THOV restriction. Variants tested in equimolar ratios include empty vector controls, wtMxA (total 2X wtMxA) (black bars, dosage control), inactive variants (red bars), H5N1 specialists (green bars), a generalist (yellow bar), or THOV specialists (purple bars). Using minireplicon assays, we also measured **(C)** H5N1 restriction and **(D)** THOV restriction of H5N1 specialist MWSRR, THOV specialist LYDRY, or their equimolar combination. All assays contained the same amount of transfected plasmid DNA. Minigenome restriction is reported as fold-change relative to an empty vector on a linear scale. For **(A)** and **(B)** we performed unpaired Welch’s t-tests between each variant and the restriction level of 2X wtMxA. For **(C)** and **(D)** we performed unpaired Welch’s t-tests between the combined specialists with each other combination. Significant differences are noted as * for p-value < 0.05. Protein expression levels were monitored for each condition by Western blotting (Supplementary Raw Images Figures 4A-D).

As expected, we found that the dominant-negative MxA variants (T103A and L542S) abrogated both H5N1 and THOV restriction compared to wtMxA (Fig. 4A-B). In contrast, the oligomerization-defective M527D variant only modestly impaired H5N1 and THOV restriction (Fig. 4A-B). Both specialist (MWSRR, SWSTR) and generalist (IYWCK) H5N1 super-restrictors continued to enhance H5N1 restriction despite their half dosage (Fig. 4A). In the THOV minireplicon, only the generalist (IYWCK) super-restrictor significantly enhanced THOV restriction at half dosage (Fig. 4B), but all other super-restrictors maintained at least wtMxA levels of restriction (Fig. 4B). Thus, in H5N1 restriction, a wtMxA allele does not interfere with super-restrictor variants. Moreover, in all cases, specialist super-restrictors do not impair wtMxA activity against the non-targeted virus, suggesting they do not act in a co-dominant or dominant-negative manner. For example, THOV-specialist super-restrictors (QFAYS, LYDRY) did not substantially lower H5N1 restriction compared to 1X wtMxA levels (Fig. 4A). Instead, we found that H5N1 specialist super-restrictors modestly enhanced THOV restriction relative to 1X wtMxA levels (Fig. 4B), despite having weak to no activity against THOV (Fig. S3B). These data indicate that MxA super-restrictors can cooperate with wtMxA to enhance antiviral restriction without impairing existing antiviral functions.

Finally, we tested whether combining an H5N1 specialist super-restrictor (MWSRR, with low THOV restriction) with a THOV specialist super-restrictor (LYDRY, with low H5N1 restriction) in heterozygous allelic combinations might achieve high restriction of both viruses. Indeed, we found this to be the case (Fig. 4C, D, Supplementary Tables S14, S15). A 1:1 mixture of the MWSRR and LYDRY variants led to a modest increase of H5N1 restriction over the half-dosage of the MWSRR variant (Fig. 4C). The same mixture also led to increased THOV restriction over the half-dosage of the LYDRY variant (Fig. 4D). These results demonstrate that the specialist super-restrictors do not act as dominant-negative variants for antiviral restriction. Instead, two specialist super-restrictor alleles in the same cell lead to simultaneously increased antiviral activity against two different viral targets.

## Discussion

Here, we demonstrate that amino acid variation within the positively selected residues in the L4 loop of MxA contains the potential to significantly enhance the restriction of the highly pathogenic avian H5N1 strain of IAV. We find that MxA super-restriction can be attributed to positive epistasis, where two different mutations in L4 combine to confer a significant gain of restriction above wtMxA levels. We also find that a critical determinant of the breadth-specificity trade-off is a single amino acid residue at position 561, with tryptophan favoring H5N1 restriction and phenylalanine or tyrosine favored in THOV restriction. Although W561 appears to prohibit THOV restriction, F561 and Y561 permit restriction of both THOV and H5N1.

Our finding that a single aromatic amino acid at position 561 determines MxA restriction activity against divergent *orthomyxoviruses* has important implications for its biochemical interactions with viral proteins and the likelihood of evolutionary transitions between different antiviral states. Several studies have pointed to the viral NP as the target of MxA antiviral action^17,19,23^. We speculate that the contrasting preference in THOV versus H5N1 super-restrictors (especially the inability of W561 to restrict THOV) results from the size or orientation of this amino acid, which might affect its interactions with THOV versus H5N1 NP. Incorporating these insights could help protein docking and modeling studies that advance our understanding of the MxA-NP interface, which is still poorly understood.

Previous evolutionary analyses have revealed that MxA genes in humans and other hominids (gorilla, chimpanzee, bonobo, orangutan) encode phenylalanine (F) at position 561. In contrast, other primate species encode a wide variety of amino acids at this critical site, including aliphatic (leucine, isoleucine, valine), sulfur-containing (cysteine, methionine), hydroxylic (serine), acidic (aspartic acid), and aromatic (tyrosine)^29^. As a result, hominoid MxA alleles are better poised than the MxA of other primates to restrict THOV and IAV^29^. Yet, despite their biochemical similarity, evolutionary transitions between F, Y, and W are not trivial. A single non-synonymous change can accomplish transitions between phenylalanine (F, encoded by codons TTT/TTC) and tyrosine (Y, encoded by codons TAT/TAC). However, transitions from phenylalanine or tyrosine to tryptophan (W, encoded by TGG) require two simultaneous transversion changes since all intermediate states (leucine, cystine, or a stop codon) would lose function against *orthomyxoviruses*^29^. Moreover, the restriction phenotype of residue 561 is reliant on epistatic interactions with other rapidly evolving residues, which in turn influence the constraints acting on residue 561.

Our analyses reveal two strategies antiviral proteins like MxA can use to bypass breadth-specificity trade-offs for orthomyxovirus restriction. First, MxA variants capable of enhanced restriction of both H5N1 and THOV can act as ‘generalist’ super-restrictors. Based on previous analyses, we anticipate this generalist super-restriction is achieved by increasing avidity to the NP proteins from both THOV and H5N1^15,30^. However, such generalist super-restrictors were rarely detected and not likely to be readily accessible in the MxA loop L4 mutational landscape.

We show that host genomes could also achieve broad enhanced restriction of divergent *orthomyxoviruses* without the requirement to evolve rare generalist super-restrictors but rather by combining two divergent alleles of MxA with distinct ‘specialist’ super-restriction activities. Since specialist super-restrictors are more common than generalist super-restrictors in MxA’s mutational landscape (Fig. 3A), we propose that heterozygous combinations of specialist super-restrictor alleles are more likely to provide host populations with a facile means to overcome the breadth-specificity tradeoffs imposed by multiple pathogenic viruses.

In theory, heterozygous MxA restriction could rely on two distinct MxA oligomers, primarily comprised of the same monomer. Because MxA restricts THOV and H5N1 in a dose-dependent manner, we would expect this to reveal a co-dominant phenotype of two alleles. However, our analyses suggest that monomers of both MxA variants intermix in heterooligomers, in which the restrictive variants can manifest their enhanced restriction even while oligomerized with less-restrictive variants. Such variants contrast with previously studied MxA variants, in which missense mutations in the MxA ‘backbone’ resulted in dominant-negative loss-of-function oligomers with wtMxA^21,35^. Such loss-of-function mutations effectively lead to loss of MxA restriction and are unlikely to propagate at high levels in populations. In contrast, gain-of-antiviral specificity variants of MxA, like those we have identified in loop L4, are expected to be positively selected, especially in the face of pathogenic viruses. They may even be co-propagated under balancing selection. Indeed, genes encoding restriction factors are often subject to diversifying and balancing selection, with heterozygote advantage maintaining multiple diverse protective alleles in host populations^37^.

Our combinatorial mutagenesis strategy focusing on positively selected residues provides a general means to elicit super-restrictor variants of restriction factors and bypass inherent breadth-versus-specificity trade-offs in antiviral restriction. The insights we have gained in the present study of MxA variants will likely apply to other restriction factors that share three critical attributes with MxA. First, like MxA, many restriction factors evolve under positive selection at their viral interaction surfaces; amino acid changes at these positively selected interfaces can confer gain of antiviral specificity and successful host restriction. Second, they can also be subject to balancing selection due to heterozygote advantages. Third, many restriction factors function as dimers or higher-order oligomers, enabling a strategy combining diverse monomeric units in oligomers to manifest a much broader antiviral restriction.

Given the immense selective pressures imposed on restriction factors like MxA, it is surprising that most mammalian genomes only encode two Mx-family proteins, one localizing to the cytoplasm and the other to the nucleus or nuclear periphery. Given similar selective pressures from pathogenic viruses, many other antiviral restriction factors have undergone dramatic expansions, like the *APOBEC3* genes in primates or the *TRIM5*-like genes in rodents or carnivore genomes^38,39^. Similarly, virus-specific alleles of the murine restriction factor *Fv-1* arose and became distributed among subspecies based on varied exposure to cocirculating viruses^40,41^. The fact that we don’t see such rampant duplication and diversification among *MxA* paralogs in mammalian genomes suggests either that two paralogs are enough to provide broad protection or (more likely) that there is some hidden cost associated with rampant Mx gene expansion in mammals, either to host fitness or due to dominant-negative interference of Mx restriction. Understanding the nature of this hidden cost would help us design better strategies to identify only slightly altered human Mx variants that could provide significantly improved antiviral protection in the face of a new IAV variant, which we will inevitably encounter in the future.

## Supporting information

Supplementary Tabes S1-S15

Western blot raw images

## Acknowledgments

We thank members of the Malik and Emerman labs for valuable discussions and Peter Dietzen and Janet Young for their comments on the manuscript. We thank Emily Hatch for the lamin antibody. This work was supported by the Genome Training Grant (5T32HG000035 to RG), Howard Hughes Medical Institute Hanna H. Gray Fellowship (GT11096/GT16732 to JLT), American Foundation for AIDS Research Mathilde Krim Fellowship in Biomedical Research (110298-71-RKHF/110537-74-RKHF, to JLT), the German Research Foundation (SFB 1160 project C01 to LG), National Institutes of Health grant (U54 AI170792 (PI: Nevan Krogan) to ME, HSM), and a Howard Hughes Medical Institute Investigator award (to HSM). Funding agencies had no role to play in the execution of the project or the decision to publish. This paper was typeset with the bioRxiv word template by @Chrelli: www.github.com/chrelli/bioRxiv-word-template

## Author contributions

Conceived the study: RAG, JLT, ME, HSM

Performed the study: RAG, DK, JLT

Project supervision: JLT, GK, LG, ME, HSM

Funding acquisition: RAG, JLT, LG, ME, HSM

Wrote the paper: RAG, ME, HSM

Edited the paper: JLT, GK, LG

## Materials and Methods

### Library Construction and Plasmid Preparation

A library of 3X-FLAG-tagged human MxA variants was designed, in which codons 540, 564, 566, and 567 were randomized by NNS mutagenesis, and codon 561 was randomly allowed to a W, F, or Y. This library was synthesized and cloned into a pQCXIP plasmid backbone. The pooled library of plasmids was transformed into NEB 5-alpha competent *E. coli* (NEB #C2987) and plated sparsely to obtain single colonies. Single colonies were inoculated into 6mL of 100 µg/mL ampicillin LB broth, grown overnight at 37°C at 250rpm. After 16-18 hours of growth, 1mL of culture was added to 1mL of 50% glycerol for storage at -80°C. The remaining 5mL were used for plasmid purification (Promega #A1223). The C-terminus of MxA, including the loop L4 region, of the purified plasmids was sequenced by Sanger sequencing using the following primer: 5-’ CGT GGT AGA GAG CTG C -3’. We cloned and sequenced ∼270 randomly selected variants from the pooled plasmid library. Of those, 194 did not contain stop codons or frameshift mutations. All variants used for follow-up assays after the initial screen were sequenced again to verify the remaining N-terminal sequence using a plas-mid-specific primer upstream of MxA (5-’ ACA CCG GGA CCG ATC CAG-3’).

### Cell Lines

HEK-293T/17 and HeLa cells were grown on treated tissue-culture plates in DMEM (Thermo Fisher #11965118) containing high-glucose and L-glutamine with 1x penicillin/streptomycin (Thermo Fisher #15140122) and 10% fetal bovine serum (Gibco #10437028). Cells were grown at 37°C, 5% CO_2_ in humidified incubators, and passaged by digestion with 0.05% trypsin-EDTA (Thermo Fisher #25300120).

### Minireplicon Assays

All minireplicon assays were performed in black, opaque, clear-bottomed 96-well plates by transfection of 50-80% confluent HEK293T/c-17 cells with minireplicon components using Mirus TransIT-293 reagent. For the H5N1 minireplicon system, 1 ng each of PB2, PB1, and PA, 0.5 ng of NP (all in a pCAGGS vector), 25 ng of pHH21-vNP-FF-Luc (firefly luciferase), 5 ng of pTK-Ren-Luc (*Renilla* luciferase), and 100 ng of pQCXIP-MxA were transfected. For the THOV minireplicon system, 4 ng each of PB2, PB1, and PA, 1 ng of NP (all in a pCAGGS vector), 20 ng of pHH21-vNP-FF-Luc (firefly luciferase), 50 ng of pTK-Ren-Luc (*Renilla* luciferase, constitutively expressed under the HSV TK promotor to serve as a transfection control), and 50 ng of pQCXIP-MxA were transfected. After 24 hours, firefly and *Renilla* luciferase luminescence were measured using the Promega Dual-Glo Luciferase Assay System. For the 96-well format, all but 20 µL of the medium was removed. To each well, 20 µL of Dual-Glo Luciferase Reagent was added for lysis and luciferase activation, incubated for 10 minutes at room temperature, and then luminescence was read on a Biotek Cytation3 plate reader. 20 µL of the Stop and Glo Reagent was added, incubated at room temperature for 10 minutes, and luminescence was re-read. Normalized minireplicon activity for each sample was calculated as the value of firefly luciferase luminescence divided by the *Renilla* luciferase luminescence. Each assay plate contained empty vector, wtMxA, and catalytically inactive MxA(T103A) controls, providing the range of control variant restriction reported in Figure 1B. Results are reported as “fold-restriction”, calculated as the average minireplicon activity in the presence of empty pQCXIP vector for the paired assay plate divided by the minireplicon activity of the experimental sample. All samples were assayed by transfection of a master mix into triplicate wells.

### Logo Plots

The DiffLogo plot was generated in R using the MotifStack and DiffLogo packages from the Bioconductor library. Frequencies of each amino acid were calculated at each site. The DiffLogo plot was generated by comparing amino acid frequencies at each site among super-restrictors to amino acid frequencies at each site across all assayed variants. The code and associated files can be found at https://github.com/rag125/h5n1_srs.

### Western Blots

We used western blot analyses to assay protein expression after plasmid transfection in HEK293T cells for 24 hours in a 24-well format. Cells were collected and lysed on ice in RIPA buffer (Invitrogen #89900). Cell debris was removed by centrifugation, after which protein concentration in all supernatants was measured using the Pierce™ BCA Protein Assay Kit and normalized to an equal concentration by diluting samples with RIPA buffer. Samples were reduced and denatured by adding Laemmli buffer (Bio-Rad #1610737) containing β-mercaptoethanol and heating to 95°C for 5 minutes. Samples were run by SDS-gel electrophoresis and blotted using the Bio-Rad Mini-PROTEAN TGX Gel system. Membranes were cut using a razor to separate MxA protein (∼76kDa) and GAPDH (∼37kDa). We used the following primary antibodies: Sigma F1804 monoclonal M2 Anti-FLAG and Genetex GTX100118 anti-GAPDH. Secondary antibodies conjugated to horseradish peroxidase (HRP) are from R&D systems (HAF007 and HAF008). HRP was detected using Supersignal West Pico Plus Chemiluminescent Substrate (Fisher #34577) on a Bio-Rad Gel Doc.

### Immunofluorescence Microscopy

HeLa cells were seeded in a 24-well plate at a density of 10^5^ cells/well. When cells were confluent about 24 hours later, they were transfected with the appropriate MxA variant equivalent to five times the volume delivered to their corresponding 96-well plates using Lipofectamine 3000 reagent (Invitrogen #L3000). Nineteen hours after transfection, cells were trypsinized and reseeded in Cellvis 24-well glass-like bottomed plates (Cellvis #P24-1.5P) at a density of 0.5-1x10^4^ cells/well. About 12 hours after reseeding, cells were washed with PBS (+Mg^2+^ +Ca^2+^) for 5 minutes, fixed in 4% paraformaldehyde for 15 minutes, rewashed, permeabilized with 0.25% Triton X-100 for five minutes, and washed again. Cells were blocked in 10% BSA at 37°C for 30 minutes. Primary antibody incubation included anti-lamin antibody (Sigma-Aldrich #L1293) at 1:500 and anti-FLAG (Sigma #F1804) at 1:2000 in 3% BSA for 1 hour at 37°C. After two PBS++ washes, cells were incubated with secondary antibodies Alexa-Fluor488 Anti-rabbit (Fisher #A21206) at 1:2000, Alexa Fluor633 Anti-mouse (Invitrogen #A21050) at 1:2000, Hoechst stain (Invitrogen #H1399) at 1:1000, and Alexa Fluor568 Phalloidin (Thermo Fisher #A12380) at 1:400 for 1 hour at 37°C. Cells were washed twice in PBS++ before imaging on a Leica DMi8 inverted microscope. Images were processed using LAS X software.

### Statistical Analyses

Statistical analyses were performed in R. The code and associated files can be found at https://github.com/rag125/h5n1_srs.

**Figure S1:**
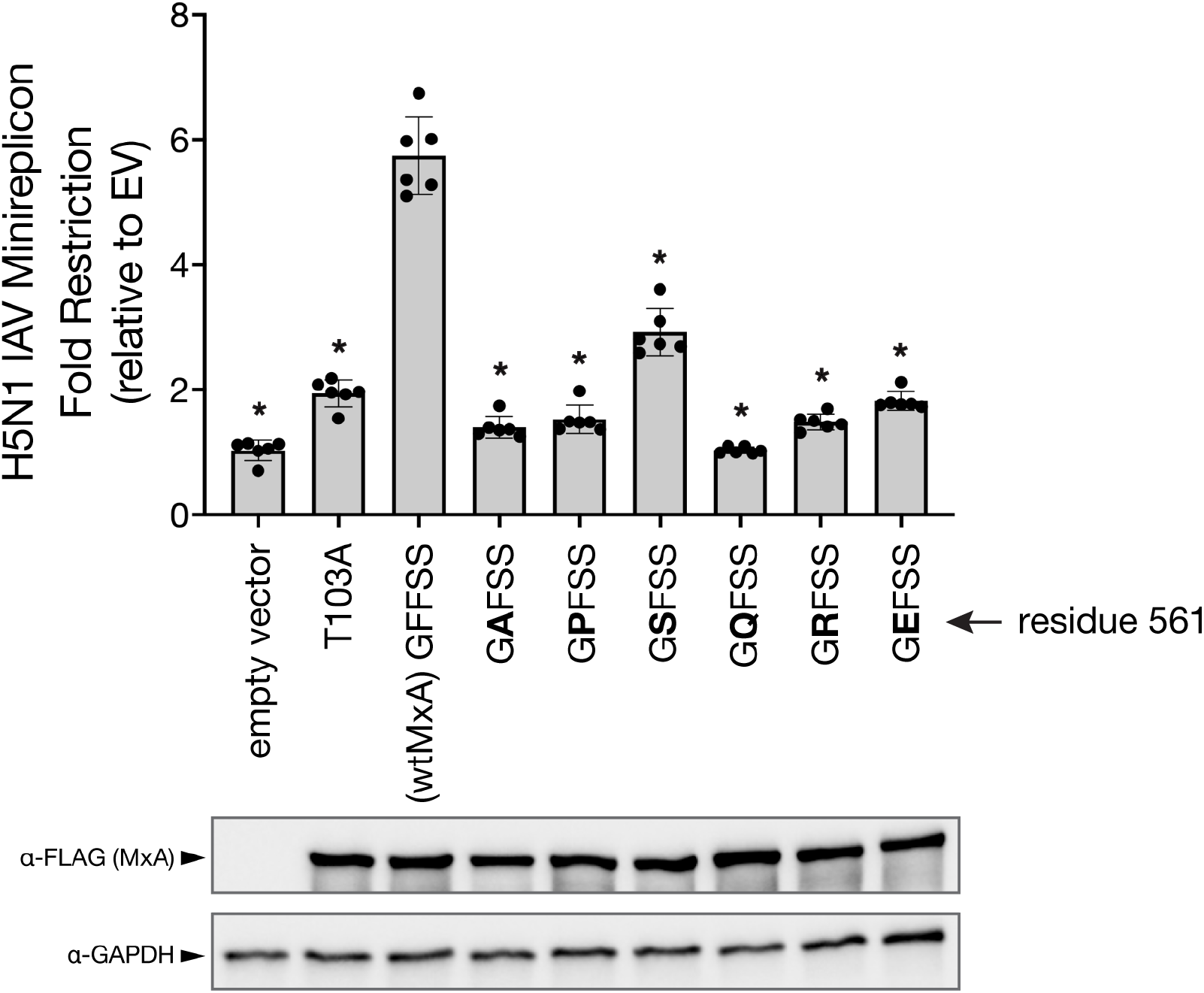
Restriction profiles of MxA variants with various non-aromatic amino acid residues at position 561. Fold restriction is reported relative to an empty vector. Each variant is labeled using amino acid identities at the five variable sites. We used unpaired Welch’s t-tests between each variant and wtMxA to evaluate statistical significance (*p-value < 0.05).

**Figure S2:**
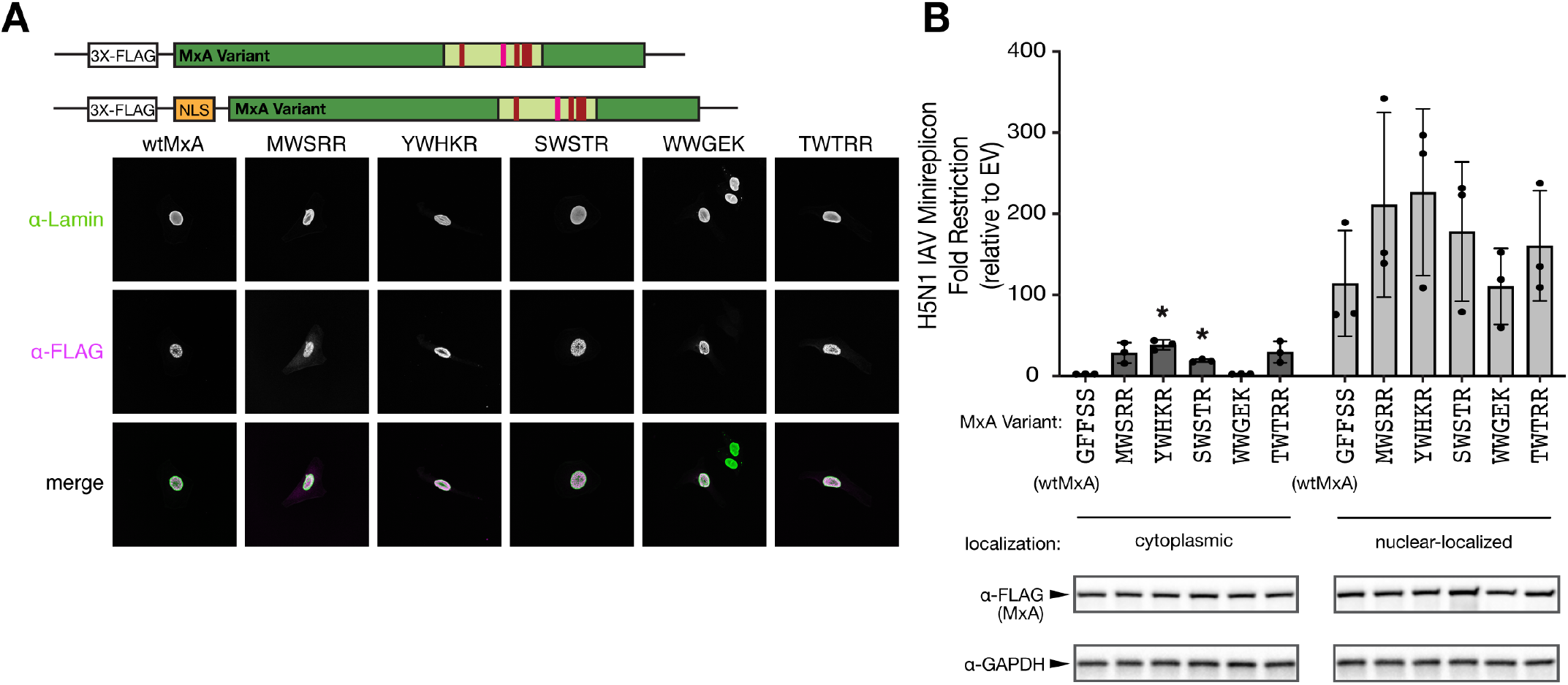
Nuclear localization further enhances H5N1 super-restriction. **(A)** The SV40 large T antigen nuclear localization signal (NLS) PKKKRKV was cloned into the N-termini of MxA variants between a 3X-FLAG tag and the MxA gene. NLS-tagged variants were transfected into HeLa cells and imaged in the same manner as described in Fig. 2A. **(B)** The five super-restrictor variants, as well as wtMxA, with and without an N-terminal NLS were assayed for their H5N1 restriction relative to an empty vector control in the minireplicon assay. Their expression levels were also tested by Western blotting. Unpaired Welch’s t-tests were performed between restriction levels of cytoplasmic variants and wtMxA as well as between NLS-tagged variants and NLS-wtMxA (* p< 0.05).

**Figure S3:**
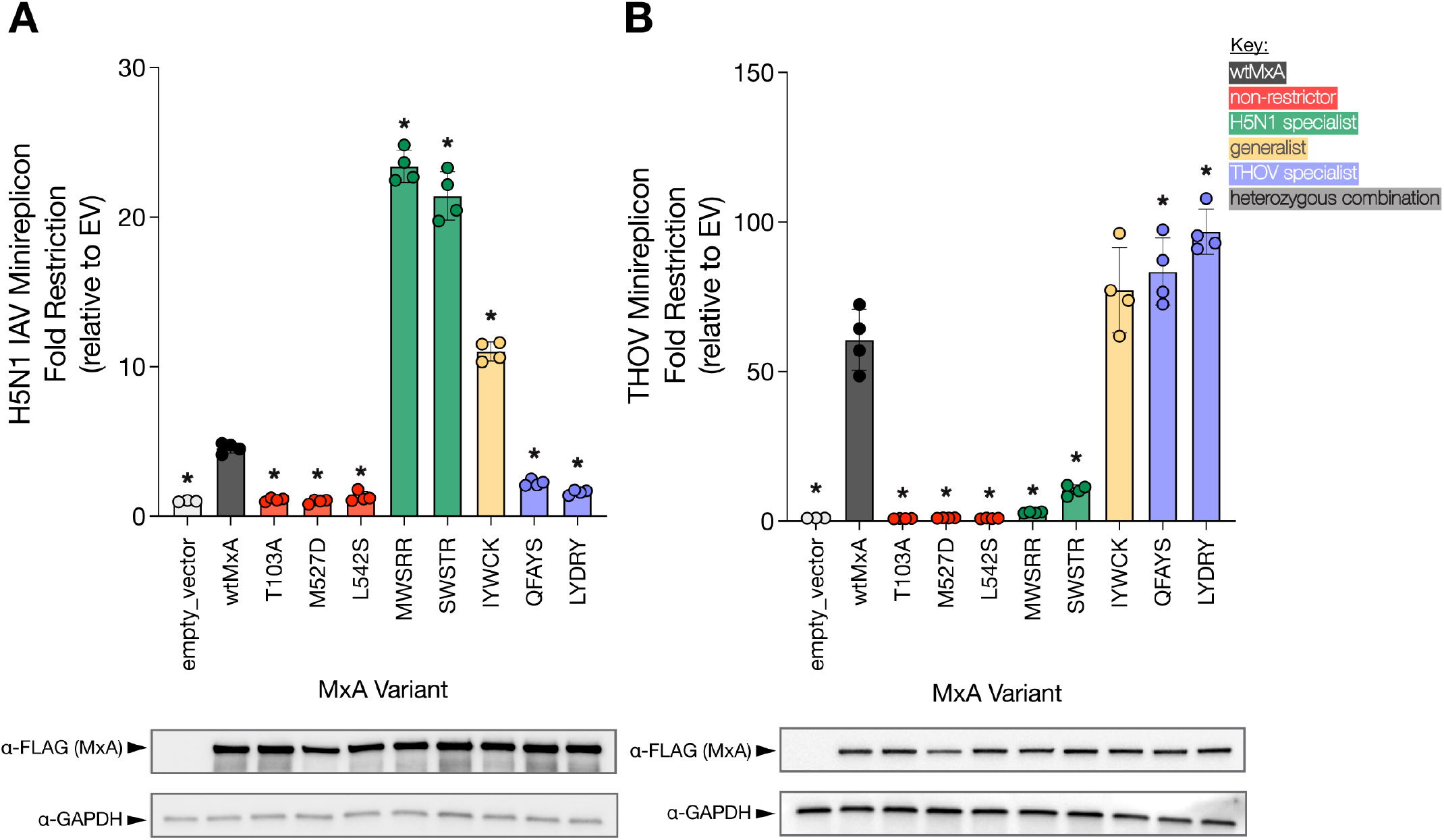
‘Generalist’ and ‘specialist’ MxA variants restriction of H5N1 and THOV. We retested H5N1 **(A)** or THOV **(B)** restriction by wtMxA, non-restricting controls, specialist, and generalist MxA variants relative to an empty vector control based on a minireplicon assay; data is represented on a linear scale. The total amount of empty vector or MxA variant per condition is 100ng per well for H5N1 assay and 50 ng per well for the THOV assay. Unpaired Welch’s t-tests were performed between restriction levels of cytoplasmic variants and wtMxA as well as between NLS-tagged variants and NLS-wtMxA (* p< 0.05).

