## Supplementary material for "Heterozygous and generalist MxA super-restrictors overcome breadth-specificity tradeoffs in antiviral restriction": Western blot raw images

### Figure 1C

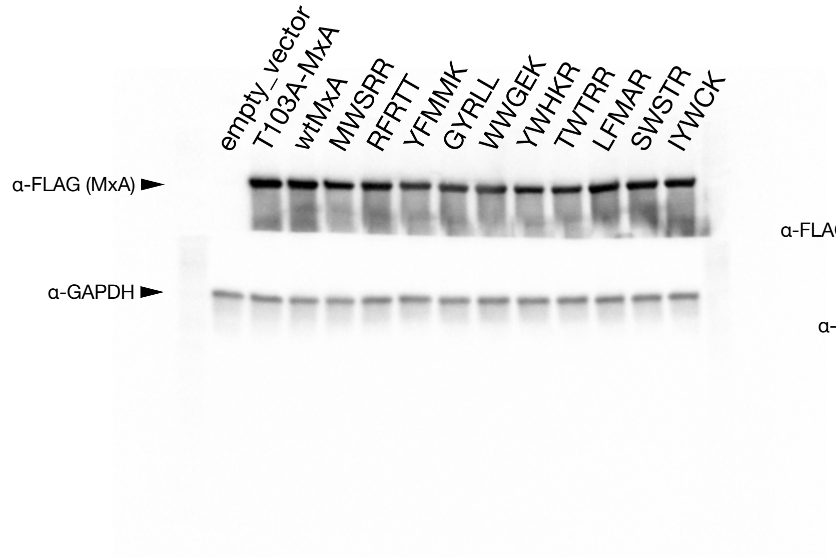


### Figure 1D

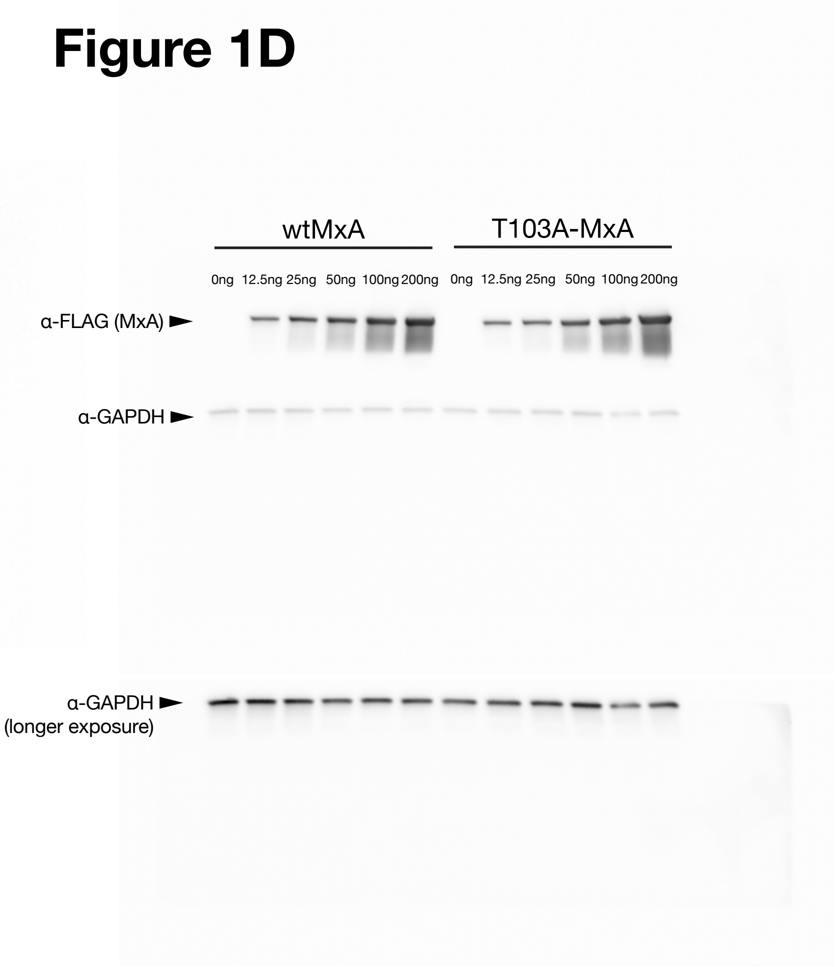


### Figure 1D

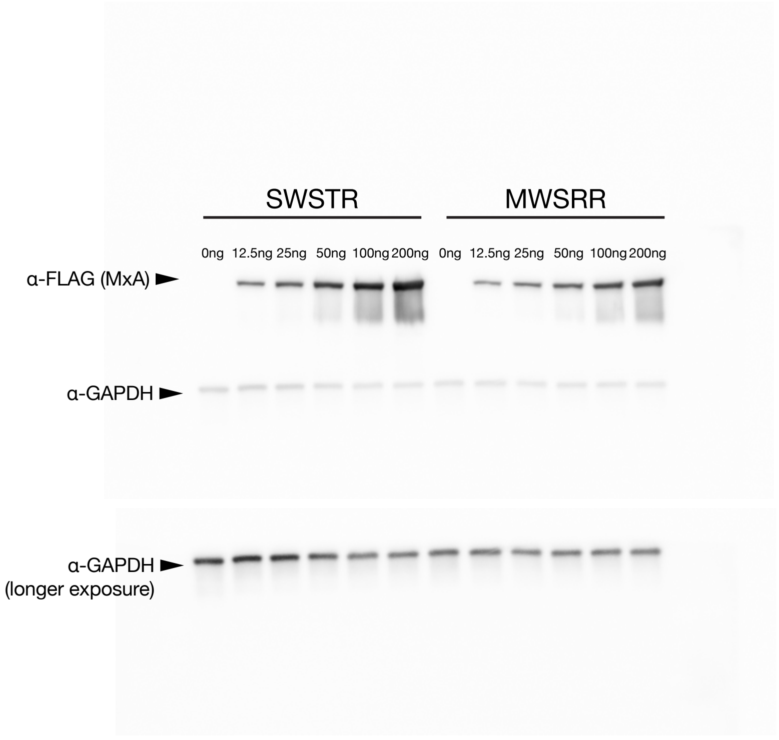


### Figure 1D

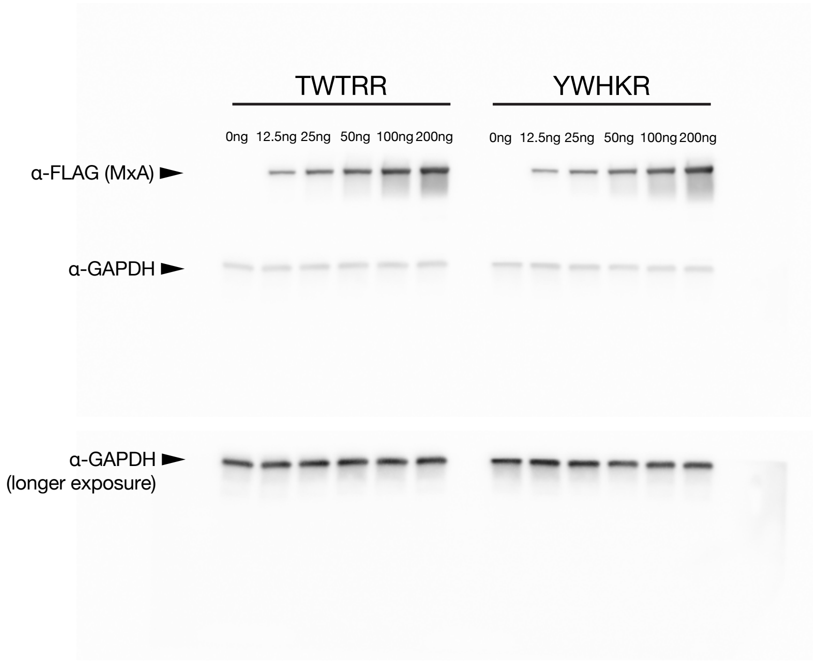

### Figure S2B

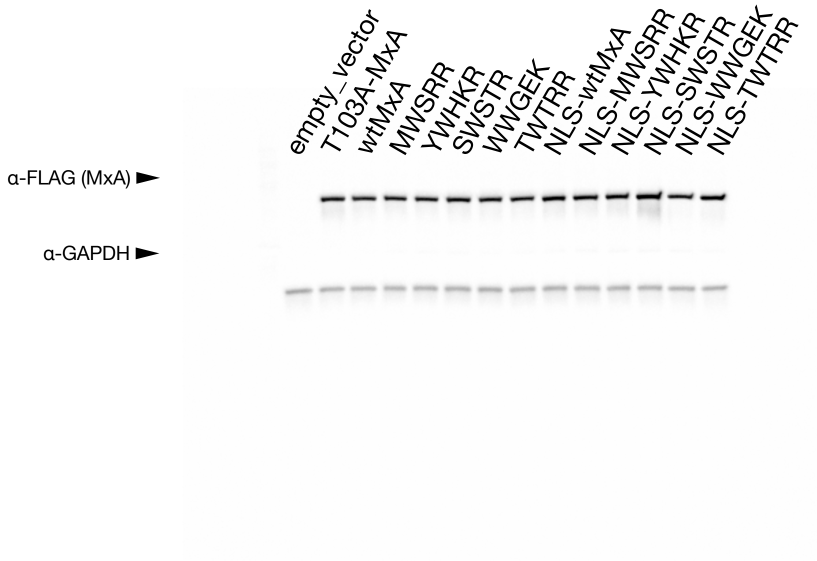


### Figure 2C - left

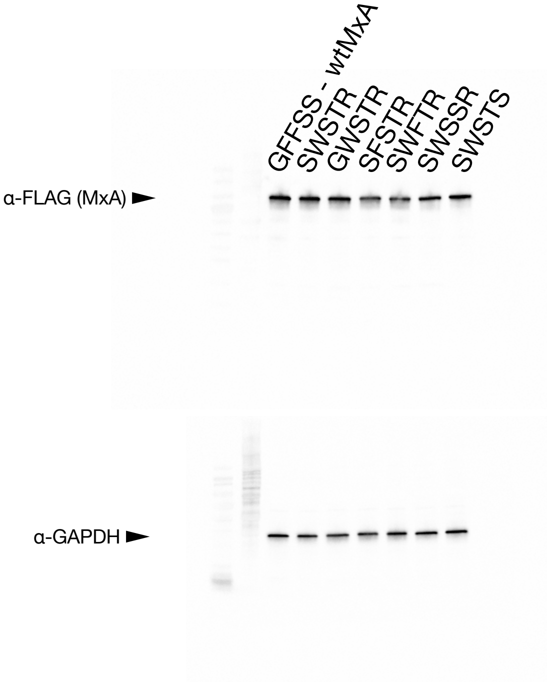


### Figure 2C - right

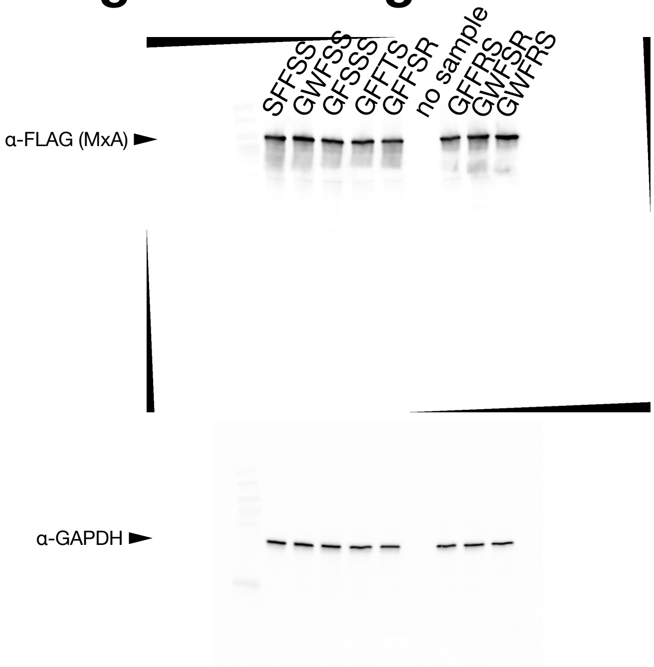


### Figure 2D

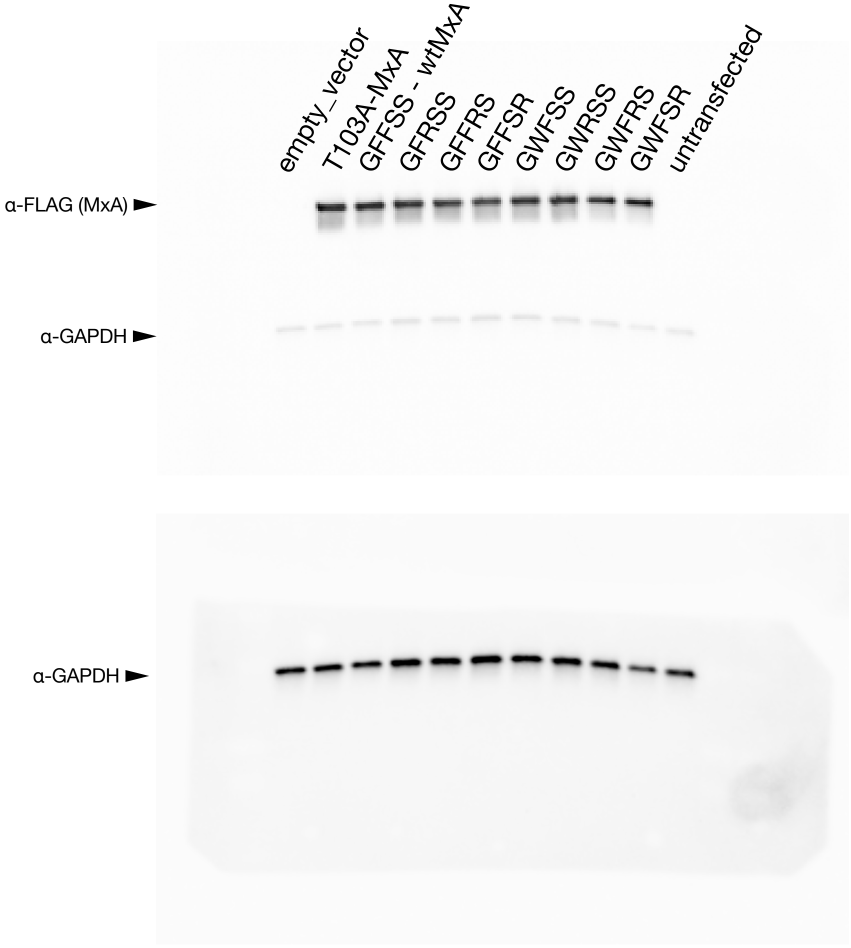

### Figure S1

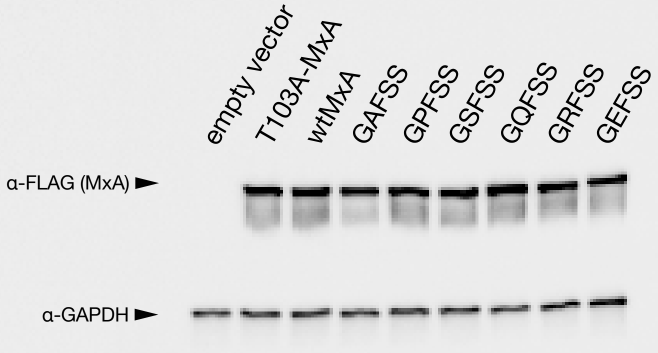


### Figure 4A

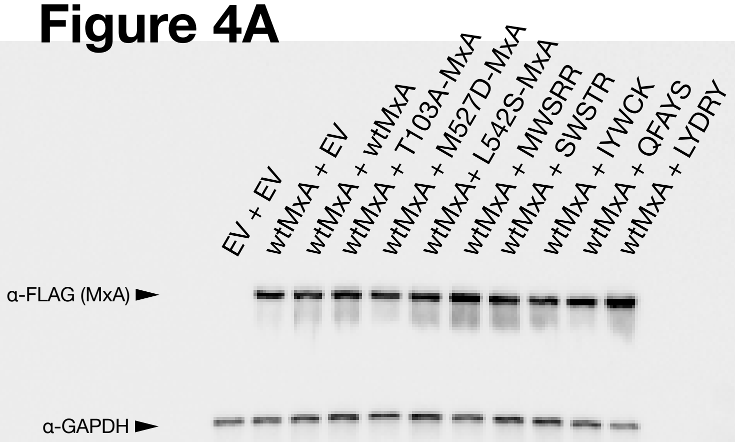


### Figure 4C

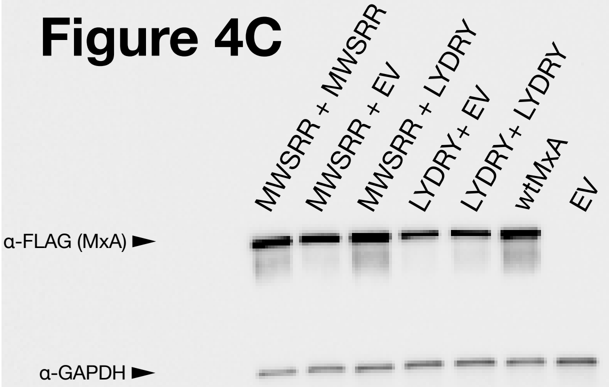


### Figure S3A

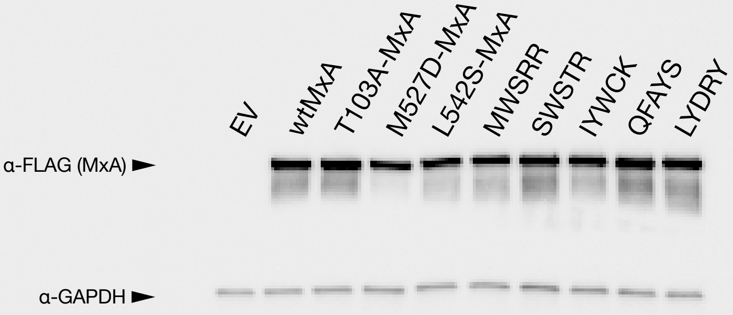

### Figure 4B

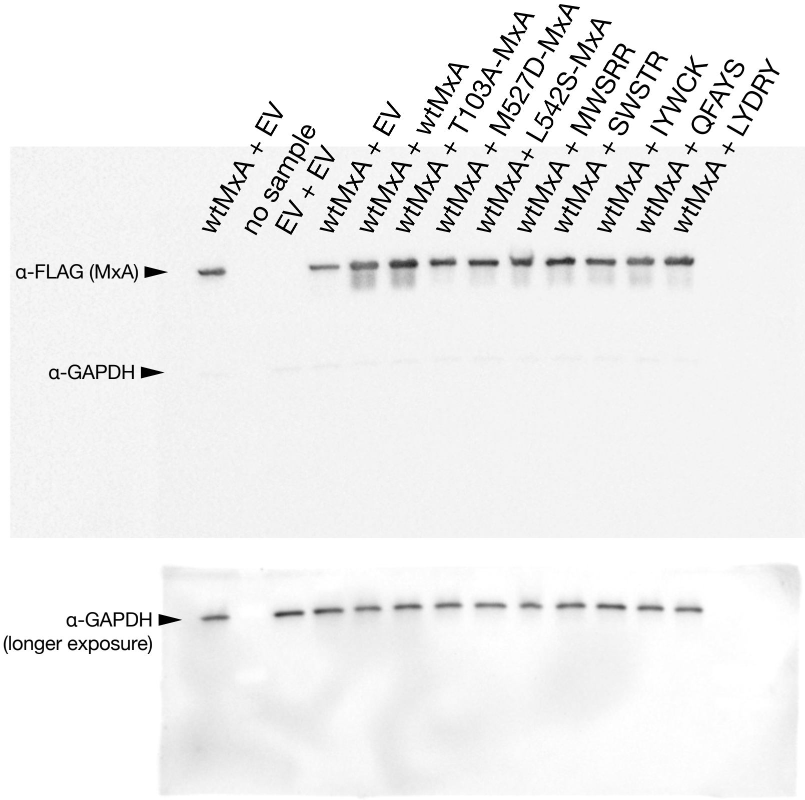


### Figure 4D

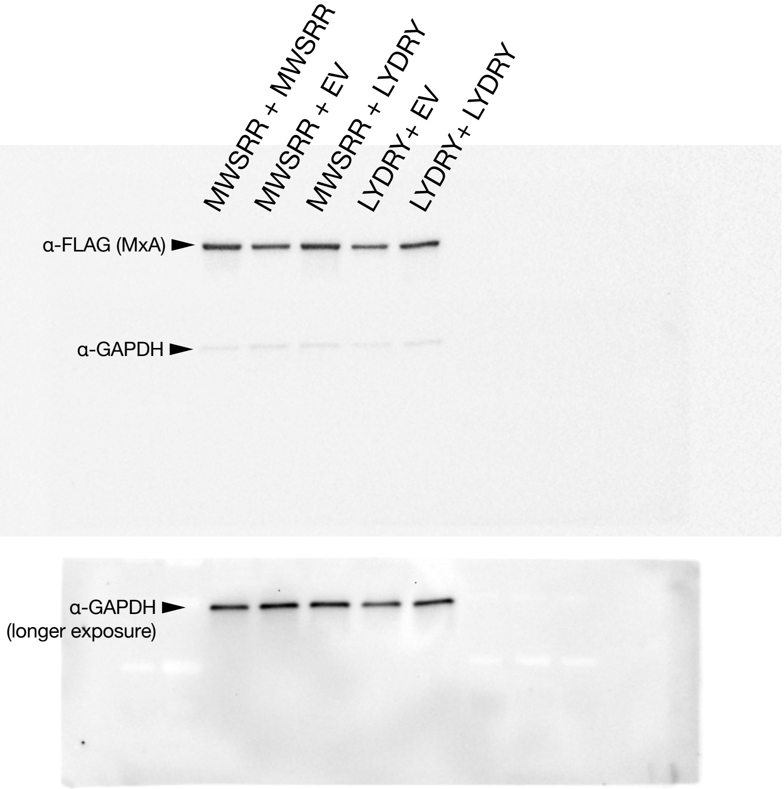


### Figure S3B

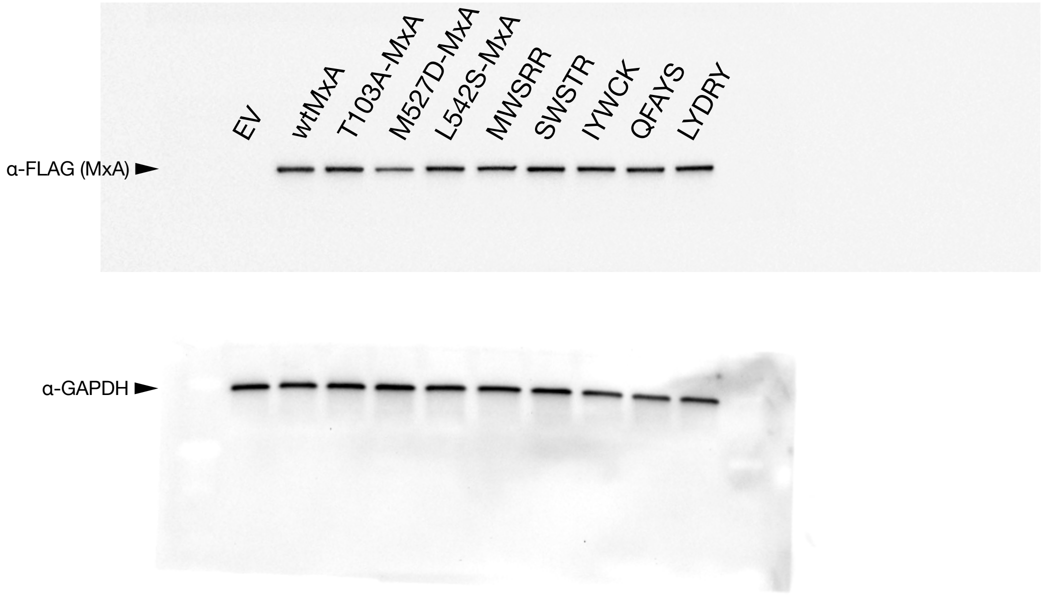
